# Enhanced BCR signalling inflicts early plasmablast and germinal centre B cell death

**DOI:** 10.1101/2020.07.29.226662

**Authors:** JC Yam-Puc, L Zhang, RA Maqueda-Alfaro, L Garcia-Ibanez, Y Zhang, J Davies, YA Senis, M Snaith, KM Toellner

## Abstract

It is still not clear how B-cell receptor (BCR) signalling intensity affects plasma cell and germinal centre (GC) B cell differentiation. We generated Cγ1^*Cre/+*^Ptpn6^*fl/fl*^ mice where SHP-1, a negative regulator of BCR signalling, is deleted rapidly after B cell activation. Although immunisation with T-dependent antigens increased BCR signalling, it led to plasma cells reduction and increased apoptosis. Dependent on the antigen, the early GC B cell response was equally reduced and apoptosis increased. At the same time, a higher proportion of GC B cells expressed cMYC, indicating increased GC B cell – Tfh cell interactions. While GC B cell numbers returned to normal at later stages, affinity maturation was suppressed in the long term. This confirms that BCR signalling not only directs affinity dependent B cell selection but also, without adequate Tfh cell help, can inflict cell death, which may be important for the maintenance of B cell tolerance.

## Introduction

Specific interaction between antigen and the B-cell receptor (BCR) is the key signal for B cell selection and activation (Niiro and Clark, 2002; Yam-Puc et al., 2018). After initial activation *in vivo*, B cells may differentiate into plasma cells (PCs) through rapid extra-follicular expansion or become germinal centre (GC) cells that will undergo BCR affinity maturation for antigen (MacLennan, 1994; MacLennan et al., 2003; Victora and Nussenzweig, 2012). GCs contribute to long-lived humoral responses by producing high-affinity antibody-forming PCs and memory B cells (MacLennan, 1994; Victora and Nussenzweig, 2012; Weisel et al., 2016). High affinity neutralising antibodies represent a crucial mechanism by which vaccines or natural infections confer sterilising immunity protecting against on re-exposure to the same pathogen (Bachmann et al., 1994; Steinhoff et al., 1995). Two major signals regulate B cell activation leading to antibody production: while signals from T helper cells have been studied intensely in recent years (Oropallo and Cerutti, 2014; Shulman et al., 2013; Victora et al., 2010), less attention has been given to the impact of BCR signalling during selection of B cells by antigen (Khalil et al., 2012; Mueller et al., 2015). The interaction between antigen and BCR controls whether activated B cells entering the GC or undergo rapid PC differentiation in extra-follicular proliferative foci. B cell clones undergoing a strong initial interaction with antigen can efficiently differentiate into extra-follicular PCs contributing to the rapid early phase of the antibody production (Paus et al., 2006). B cells expressing a wide range of BCR affinities become pre-GC B cells after T-B interaction (Dal Porto et al., 2002; Schwickert et al., 2011; Victora et al., 2010). Higher affinity BCRs can induce stronger signal transduction than lower affinity ones (Kouskoff et al., 1998). BCR occupancy is a product of BCR affinity and antigen concentration, and concentration of free antigen can be limited by antibody feedback (Toellner et al., 2018). The effect of all of this on cell fate decisions during B cell differentiation merits more attention.

The Src homology 2 (SH2) domain-containing protein-tyrosine phosphatase (PTP)-1 (SHP-1), encoded by the Ptpn6 gene, negatively regulates BCR signalling primarily via its binding to the immunoreceptor tyrosine-based inhibitory motif (ITIM)-containing receptors CD72, CD22, FcγRIIB and paired Ig-like receptor (PIR)-B (Adachi et al., 2001; D’Ambrosio et al., 1995; Maeda et al., 1998; Nitschke and Tsubata, 2004). SHP-1 is expressed and constitutively activated in all B cells and its specific deletion on B cells results in systemic autoimmunity (Pao et al., 2007). SHP-1 is highly expressed and activated in GC B cells, suggesting that BCR signalling is negatively regulated during differentiation of GC B cells (Khalil et al., 2012). While BCR signalling has been shown to be absent in dark zone (DZ) GC B cells (Stewart et al., 2018), there is more signal transduction in light zone (LZ) B cells competing for selection signals through affinity-dependent activation of their BCR (Mueller et al., 2015).

In order to test how BCR signalling inhibition by SHP-1 affects antigen-induced B cell differentiation, we generated Cγ1^*Cre/+*^Ptpn6^*fl/fl*^ mice, in which the T-dependent B cell activation induces SHP-1 deletion in most B cells (Roco et al., 2019). Most induction of immunoglobulin class switch recombination (CSR) happens during the initial phase of cognate T cell - B cell interaction before GCs are formed (Marshall et al., 2011; Roco et al., 2019; Toellner et al., 1998; Zhang et al., 2016). Using this system, we show that Cγ1^*Cre/+*^Ptpn6^*fl/fl*^ B cells exhibit stronger BCR signalling. Paradoxically this leads to a smaller extra-follicular IgG1^+^ PC response and to death of GC B cells, resulting in reduced affinity maturation in the GC.

## Results

### Increased apoptosis in extra-follicular plasma cells of Cγ1^*Cre/+*^Ptpn6^*fl/fl*^ mice

B cells binding antigen with higher affinity are more likely to differentiate into extra-follicular PCs (O’Connor et al., 2006; Paus et al., 2006). To test whether deletion of the negative regulator of BCR signalling, SHP-1, affects the early extra-follicular PC response to immunisation, Cγ1^*Cre/+*^Ptpn6^*fl/wt*^ and Cγ1^*Cre/+*^Ptpn6^*fl/fl*^, in the following abbreviated as Shp1^*fl/wt*^ and Shp1^*fl/fl*^ mice, were immunised with SRBCs i.v.. The Cγ1^*Cre*^ allele reports expression of IgG1 germline transcripts (Casola et al., 2006). These are strongly induced after the initial interaction of B cells with T helper cells before PCs or GCs appear (Marshall et al., 2011; Roco et al., 2019; Zhang et al., 2018). This should lead to efficient deletion of SHP-1 in extra-follicular PCs and GC founder B cells. Spleens were analysed 5 days post immunisation, when the extra-follicular PC response peaks and early GCs have formed (Zhang et al., 2018).

Against expectation, flow cytometry showed that PC numbers were reduced by 50% in Shp1^*fl/fl*^ mice (Fig. 1A). This primarily affected IgG1 switched PCs, while non-switched IgM PCs developed in similar numbers as in Shp1^*fl/wt*^ control animals (Fig. 1B). Immunohistology, using IRF4 as a marker for PCs, showed reduced PC foci in the splenic red pulp, primarily in the IgG1-switched PCs of Shp1^*fl/fl*^ mice (Fig. 1C). PCs emerging from GCs at the GC-T zone interface (GTI) (Zhang et al., 2018) were unaffected at this point (Supp. Fig. 1). Taken together, these data indicate that increased BCR signalling after initial B cell activation inhibits extra-follicular PC differentiation.

**Figure 1.**
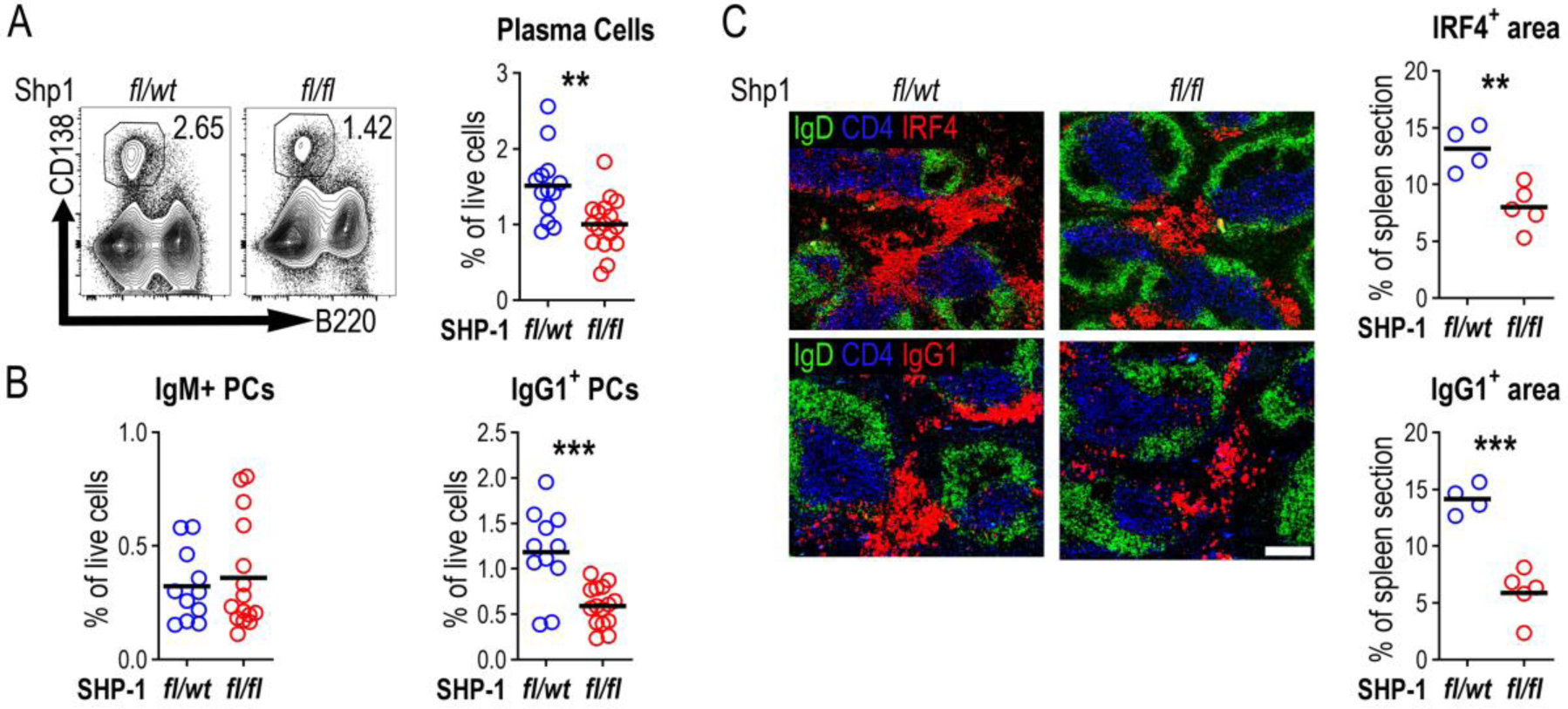
Plasma cells are reduced in Cγ1^Cre/+^Ptpn6^*fl/fl*^ mice post SRBCs immunisation. Mice spleens were analysed 5 days post immunisation with SRBCs **A**. Representative contour plots gating PCs (lymphocytes/singlets/live/B220^-^CD138^+^). Right: Summary data (% of live cells; data combined from three independent experiments. WT, n=11; Shp1, n=17). **B**. Percentage of IgM^+^ and IgG1^+^ PCs (% of live cells; data combined from three independent experiments. WT, n=11; Shp1, n=17) **C**. Splenic sections from Shp1^*fl/wt*^ (*fl/wt*) and Shp1^*fl/fl*^ (*fl/fl*) mice staining B cell follicles (IgD, green), T cell zone (CD4, blue) and PCs (top, IRF4 in red), or IgG1+ cells (bottom, IgG1 in red), scale bar 200 μm. Positive area of IRF4 and IgG1 was calculated as percentage of total splenic area. Data are representative of one of two independent experiments, WT, n=4; Shp1, n=5. Horizontal lines indicate the mean. *P < 0.05, **P < 0.01 (parametric two-tailed unpaired t-test).

Hyper-activation of B cells through BCR signalling can lead to programmed cell death (Akkaya et al., 2018; Parry et al., 1994; Tsubata et al., 1994a; Tsubata et al., 1994b; Watanabe et al., 1998). In order to test whether cell death was responsible for the smaller extra-follicular PC response, apoptotic cells were detected using Annexin V and 7-AAD staining. This showed an increase in the proportion of apoptotic PCs (Annexin V^+ve^ and 7-AAD^+ve^) in Shp1^*fl/fl*^ mice (Fig. 2A). Furthermore, the expression of active caspase-3 on different isotypes of PCs showed that both IgG1^+^ and to some extent IgM^+^ PCs of Shp1^*fl/fl*^ animals were more likely to express active caspase-3 (Fig. 2B). Immunohistology confirmed an increase in active caspase-3^+^ cells in the IRF-4^+^ extra-follicular splenic foci of Shp1^*fl/fl*^ mice (Fig. 2C). While the increase in apoptosis appears to be relatively small (+20%), it is worth mentioning that cells at this late stage of apoptosis are rapidly removed (Hanayama et al., 2004). Therefore, this likely underestimates the amount of apoptosis at this specific stage of the response, indicating that an inappropriate increase in BCR signalling can negatively affect extra-follicular PC generation through increased cell death.

**Figure 2.**
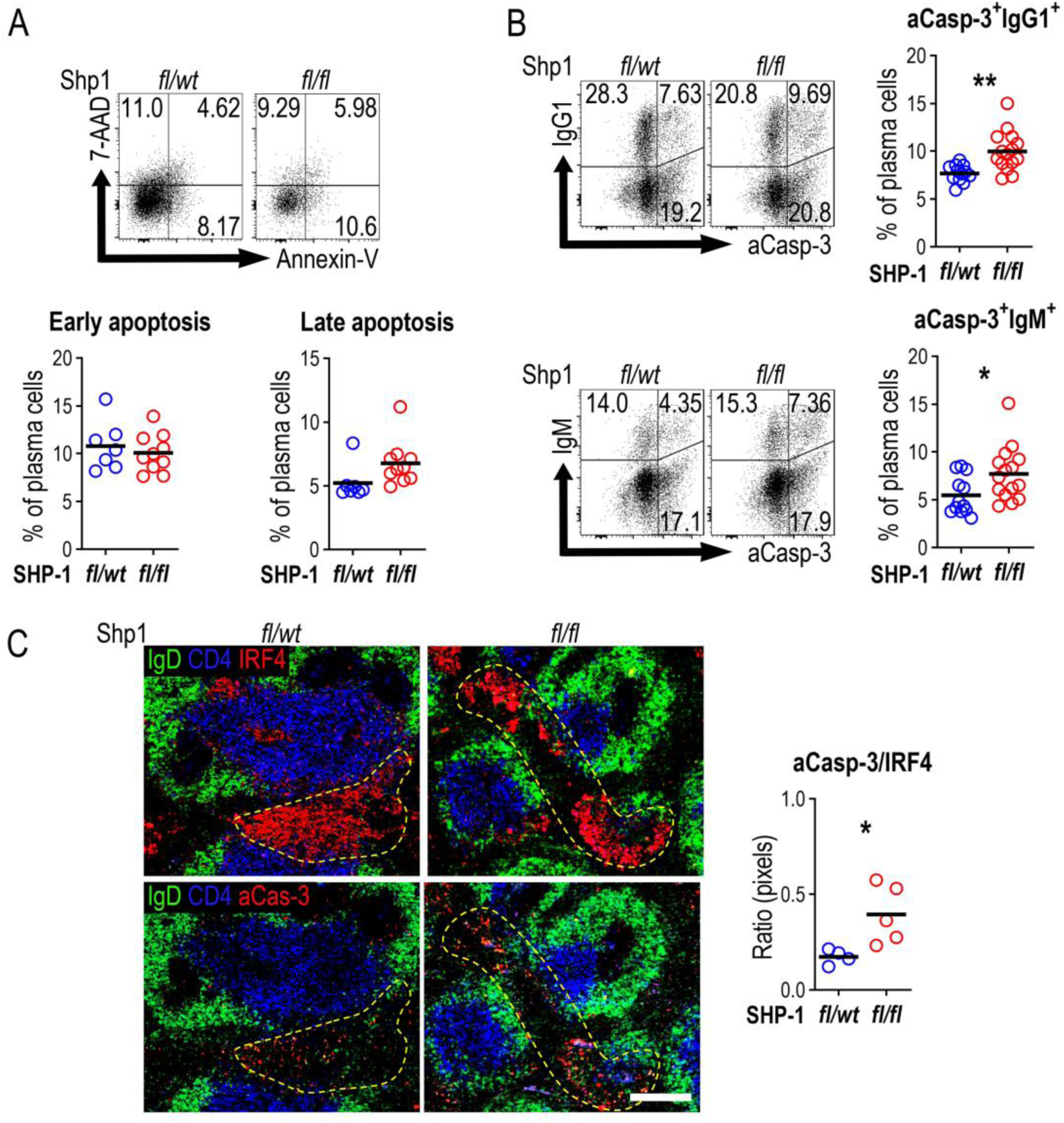
Plasma cell apoptosis is increased in SRBC immunised Cγ1^Cre/+^Ptpn6^*fl/fl*^ mice. Apoptosis rate on PCs was analysed 5 days post SRBCs immunisation in Cγ1^*Cre/+*^Ptpn6^*fl/wt*^ (*fl/wt*) and Cγ1Cre/+Ptpn6^*fl/fl*^ (*fl/fl*) mice. **A**. Representative dot plots show apoptosis rate based on the binding of Annexin V and the dead cell dye 7-AAD (pregated on PCs; top panel). Annexin V^+^ 7-AAD^-^ cells were considered as early apoptotic cells and Annexin V^+^ 7-AAD^+^ cells as late apoptotic cells. Summary data (bottom panel; % of plasma cells; results are combined from two independent experiments. WT, n=7; Shp1, n=10. **B**. Active caspase-3 expression on IgG1^+^ or IgM^+^ PCs. Graphs on the right show summary of data as percentage of active caspase-3^+^ cells (% of plasma cells; results are combined from three independent experiments. WT, n=11; Shp1, n=17). **C**. Spleen sections from Shp1^*fl/wt*^ (*fl/wt*) and Shp1^*fl/fl*^ (*fl/fl*) mice staining for B cell follicles (IgD, green), T cell zone (CD4, blue) and PCs (IRF4 in red) in the top, or active caspase-3^+^ cells (Caspase-3 in red) in the bottom. Ratio of active caspase 3^+^ pixel / IRF4^+^ pixel, representative of one of two independent experiments. WT, n=4; Shp1, n=5. Horizontal lines indicate the mean. *P < 0.05, ***P < 0.01 (parametric two-tailed unpaired t-test).

### SRBC induced GCs of Shp1^*fl/fl*^ mice are largely unaffected

In established GCs, BCR signalling is limited by SHP-1 hyper-phosphorylation, and this is important for GC maintenance (Khalil et al., 2012). In order to test how SHP-1 deletion, starting from the earliest stages of GC development, affects the GC response we followed GC B cell differentiation in Cγ1^*Cre/+*^Ptpn6^*fl/fl*^ mice 5 d after SRBC immunisation. Surprisingly, at this early stage there was no significant change in the number of GC B cells in Shp1^*fl/fl*^ mice (Fig. 3A). Flow cytometry confirmed a reduction of SHP-1 staining intensity in all GC B cells (Supp. Fig. 2A), indicating most GC B cells successfully deleted the gene. The increased in SYK phosphorylation seen in GC B cells in Shp1^*fl/fl*^ mice confirmed that SHP-1 deletion does increase BCR signalling in this system (Fig. 3B). In contrast to what was seen in extra-follicular PCs, cell death in GC B cells, evaluated by flow cytometric analysis of Annexin V / 7-AAD and active caspase 3 staining, was not increased at this stage (Fig 3C).

**Figure 3.**
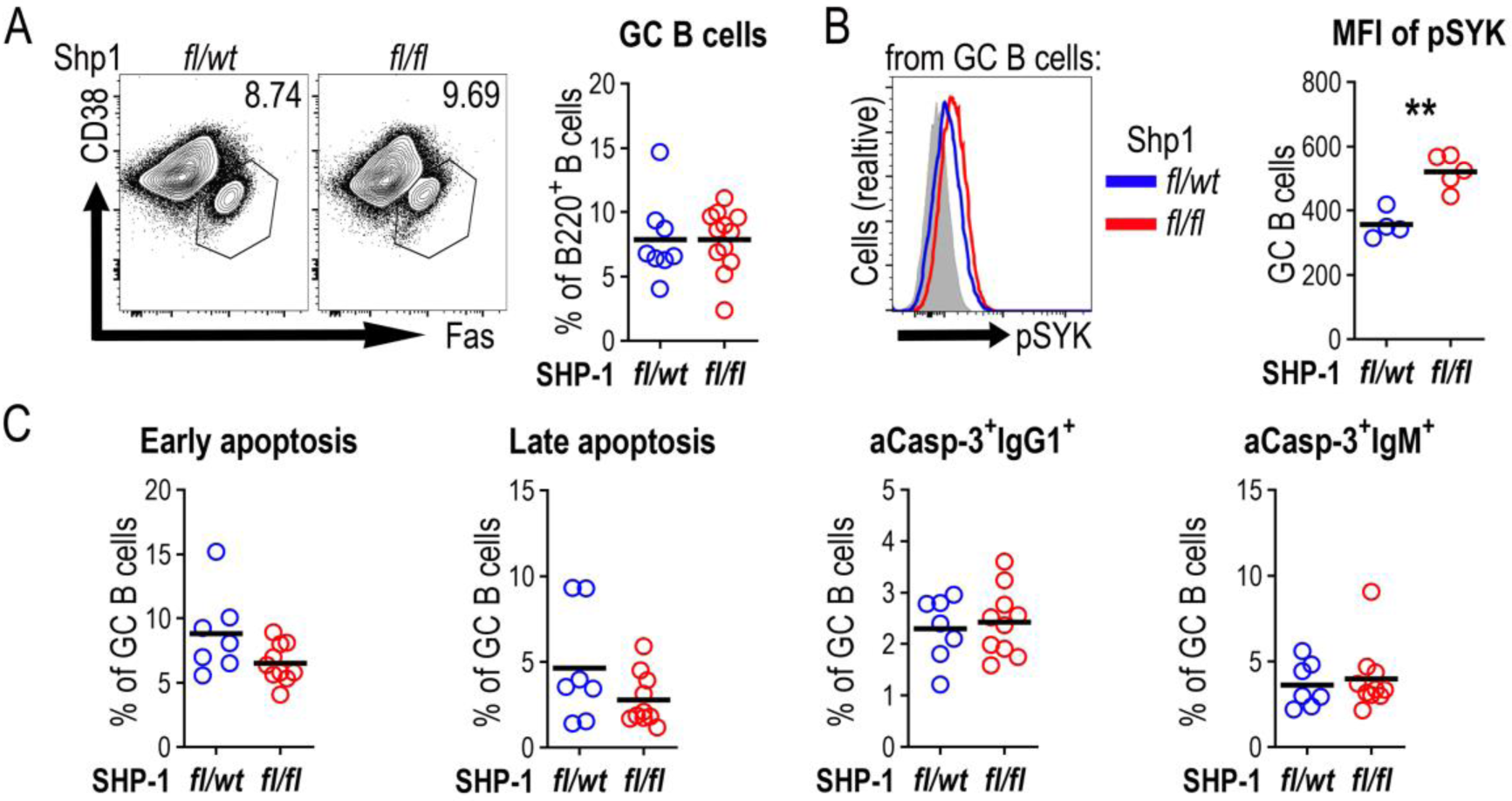
SRBC induced GCs of Cγ1^Cre/+^Ptpn6^*fl/fl*^ mice are largely unaffected. Germinal centre response was analysed 5 days post SRBC immunisation of Cγ1^*Cre/+*^Ptpn6^*fl/wt*^ (*fl/wt*) and Cγ1^*Cre/+*^Ptpn6^*fl/fl*^ (*fl/fl*) mice. **A**. Representative contour plots gating GC B cells from spleen (lymphocytes/singlets/live/CD138^-^B220^+^CD38^-^Fas^+^). Right panel shows summary of data (% of B220^+^ B cells; data are combined from two independent experiments. WT, n=8; Shp1, n=12). **B**. pSYK expression in GC B cells. Right panel shows summary data (median fluorescence intensity (MFI) of GC B cells; results are representative of two independent experiments. WT, n=4; Shp1, n=5). **C**. Apoptosis rate based on the binding of Annexin V and 7-AAD, active caspase-3 in IgG1^+^ or IgM^+^ cells as in Figure 2. (% of GC B cells; data are combined from two independent experiments. WT, n=7; Shp1, n=10) horizontal lines indicate the mean. *P < 0.05 (parametric two-tailed unpaired t-test).

### Germinal centre B cell responses and affinity maturation to NP-CGG are impaired in Cγ1^*Cre/+*^Ptpn6^*fl/fl*^ mice

While SRBC immunisation rapidly induces B cell activation and differentiation (Zhang et al., 2018), it also has a substantial T independent component and therefore is not necessarily the strongest inducer of IgG1 germline transcription. Protein antigens in alum induce strong Th2 type B cell activation, IL-4 expression in T cells, and rapid extra-follicular PCs as well as GC differentiation in draining lymph nodes (Toellner et al., 1998).

Eight days post s.c. immunisation with NP-CGG, Shp1^*fl/fl*^ draining lymph nodes showed a similar level of PC reduction than in Shp1^*fl/fl*^ spleens after SRBC immunisation. GC B cells were also reduced (Fig. 4A, B). Again, SHP-1 deletion (Supp. Fig. 2B) resulted in increased BCR signalling, as detected by increased SYK phosphorylation (Fig. 4C). Further, cMYC expression, induced after GC B cell interaction with Tfh cells (Calado et al., 2012; Dominguez-Sola et al., 2012; Luo et al., 2018), was increased in Shp1^*fl/fl*^ GC B cells (Fig. 4D), suggesting also more efficient T-dependent B cell activation. Despite this, apoptosis in GC B cells was increased (Fig. 4E). Deletion of C-terminal Src kinase (CSK), another downstream inhibitor of BCR signalling, led to a similar reduction of the GC B cell response in Cγ1^Cre/+^CSK^*fl/fl*^ mice after NP-CGG immunisation (Supp. Fig. 3).

**Figure 4.**
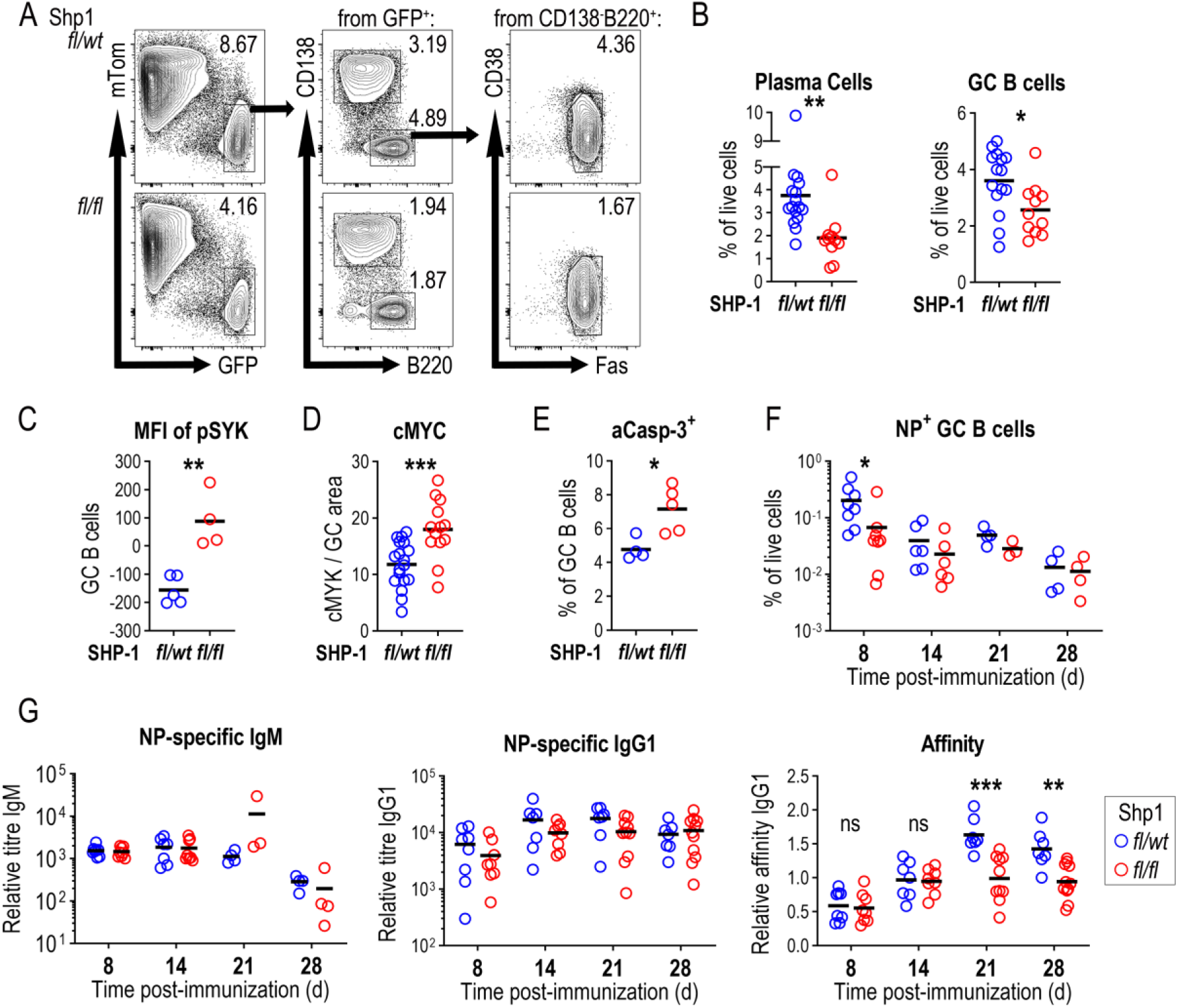
Germinal centre B cell responses and affinity maturation to NP-CGG are impaired in Cγ1^*Cre/+*^Ptpn6^*fl/fl*^ mice. **A)** Germinal centre B cell responses in popliteal lymph nodes (PLN) d8 post NP-CGG immunisation of Cγ1^*Cre/+*^Ptpn6^*fl/wt*^ (*fl/wt*) mTmG and Cγ1^*Cre/+*^Ptpn6^*fl/fl*^ (*fl/fl*) mTmG mice. Sequential gating strategy for identification of PCs (lymphocytes/singlets/live/tomato^-^GFP^+^B220^-^CD138^+^) and GC B cells (lymphocytes/singlets/live/tomato^-^GFP^+^CD138^-^B220^+^CD38^-^Fas^+^). **B**. Summary data of PCs and GC cells (% of live cells; three independent experiments; WT, n=15; Shp1, n=11). **C**. SYK phosphorylation in GC B cells from PLN 8 days post NP-CGG immunisation (MFI on GC B cells; representative results from one of two independent experiments. WT, n=5; Shp1, n=4). **D**. Summary data of relative percentage of GC cMYC expression measured from stained PLN sections 8 days post NP-CGG immunisation. (Each dot represents a different GC. Data combined from two independent experiments. WT, n=8; Shp1, n=10). **E**. Active caspase-3 on GC B cells from PLNs 8 days post NP-CGG immunisation (% of GC B cells; results are representative of one of two independent experiments. WT, n=5; Shp1, n=4). **F**. Splenic NP-specific GC B cells at different time points after immunisation (% of live cells; results are from one to two independent experiments at each time-point. WT, n=4-8; Shp1, n=3-8). **G**. Serum antibody titres for NP-specific IgM (left panel, WT, n=4-8; Shp1, n=3-8), IgG1 (middle panel, WT, n=7-8; Shp1, n=8-11), and relative IgG1 NP affinity (right panel, WT, n=7-8; Shp1, n=8-11) at different time points post immunisation. (Results are from one to two independent experiments at each time-point); horizontal lines indicate the mean. *P < 0.05, **P < 0.01, ***P < 0.001, ****P < 0.0001 (parametric two-tailed unpaired t-test and two-way ANOVA).

In order to test the effects of SHP-1 deletion on later stages of the response to TD antigens, the splenic GC response and affinity maturation were monitored after intraperitoneal immunisation of Shp1^*fl/fl*^ and Shp1^*fl/wt*^ mice with NP-CGG. Similar to the early GC response in lymph nodes, increased BCR signalling led to a reduced splenic GC response by day 8 after immunisation, but this effect was lost at later stages of the response (Fig. 4F). As expected, antigen-specific IgM production was unaffected, and NP-specific IgG1 was marginally reduced at the earliest stages of the response. At later stages, however, there was no difference in antibody titre and after 14 d after immunization Shp1^*fl/fl*^ mice did not further increase affinity of NP-specific serum IgG (Fig. 4G), showing that despite the normalization in cell numbers, there is a long-term effect on the efficiency of affinity dependent B cell selection.

## Discussion

During development B cells are strictly selected for the continuous expression of a functional BCR and the loss of its expression leads to rapid cell death (Kraus et al., 2004; Lam et al., 1997). Altering the expression of downstream signalling molecules changes the BCR signalling strength which in turn affects B cell development, (Cariappa et al., 2001; Nguyen et al., 2017; Pao et al., 2007; Tsiantoulas et al., 2017). For example, enhanced BCR signalling due to specific deletion of SHP-1 in all B cells leads to B1a B cell subset expansion (Pao et al., 2007). The effect of altering BCR signalling strength also depends on the phase of development. Enhanced BCR signalling in transitional B cells favours follicular B cell development (Cariappa et al., 2001). Few studies have tested the effects of artificially enhanced BCR signalling in mature B cells that had undergone normal B cell development (Davidzohn et al., 2020; Li et al., 2014). The model presented here allows normal B cell development and increased BCR signalling by deletion of SHP-1 only after mature and naïve B cells are activated by signals that may induce class-switching to IgG1.

In mature naïve B cells, antigen binding induces a cascade of phosphorylation involving the accessibility of immunoreceptor tyrosine-based activation motifs (ITAMs) on Igα (CD79A) and Igβ (CD79B), leading to activation of key molecules to finally activate transcription factors and prompt an immune response (Buchner and Muschen, 2014). SHP-1 inhibits BCR signalling through SYK (Adachi et al., 2001), and activated B cells in the current model show clear signs of pSYK overexpression after B cell activation.

After antigen mediated BCR stimulation and T cell help, B cells may differentiate into extra-follicular PCs (MacLennan et al., 2003) or become GC precursor cells to start GC reactions (MacLennan, 1994; Victora and Nussenzweig, 2012). The exact mechanisms which control the differentiation of B cells into extra-follicular PCs or GCs is still controversial and under scrutiny. It has been shown that B cells experiencing a strong initial interaction with antigen more efficiently differentiate into extra-follicular PCs (Paus et al., 2006). Here, we show that higher signalling through the BCR affects PC differentiation in unexpected ways. Cγ1 germline transcripts are induced during the initial B cell activation prior to extra-follicular or GC B cell differentiation (Marshall et al., 2011; Roco et al., 2019). Therefore, early B blasts differentiating into extra-follicular plasmablasts would be the first to encounter Cre mediated deletion of SHP-1 and increased BCR signalling. As increased BCR signalling should enhance extra-follicular PC differentiation (Paus et al., 2006), it was surprising to see reduced numbers of extra-follicular plasmablasts. Due to the low number of B cells activated in a non-BCR transgenic animal it was not possible to test whether the number of B cells initially activated to enter plasmablast differentiation was changed. Stronger BCR mediated activation may well have led to larger numbers of B cells entering plasmablast differentiation, however, stronger activation in the absence of co-stimulation from T cells can also induce activation-induced cell death (Akkaya et al., 2018; Parry et al., 1994; Tsubata et al., 1994a; Tsubata et al., 1994b; Watanabe et al., 1998). This would suggest that after activation, SHP-1-deficient B cells are not maintained since they do not receive timely co-stimulatory signals needed for full activation (Akkaya et al., 2018). These results are in line with data from an earlier study (Li et al., 2014) that showed a modest reduction in PC production in the response to primary immunisation with TD antigens in mice where Ptpn6 is deleted by Cre expressed under the control of the Aicda promotor (Li et al., 2014). Aicda is also induced during primary B cell activation before GCs form, however, its expression starts slightly later and at lower levels than Cγ1 germline transcripts (Roco et al., 2019), which may explain the more subtle changes.

The effect of SHP-1 deletion on GC size is only transient which could be due to the expansion of a minority of cells with incomplete deletion. The longer-term change in affinity maturation, however, makes it more likely that the complex balance between affinity dependent GC B cell selection, proliferation, output, and death reaches a new equilibrium, filling GC B cell niches to normal occupancy levels. This may explain differences seen to an earlier study, where tamoxifen-induced deletion of SHP-1 during the peak of the GC response resulted in a rapid loss of GC B cells within a short period (Khalil et al., 2012).

Although the effect on the size of the GC compartment in NP-CGG immunised mice was only transient, the higher pSYK levels clearly indicate considerably increased signal transduction in GC B cells. pSYK levels were also increased in GC B cells induced by SRBC immunization, although there was less obvious effect on GC size. This may be explainable by the fact that the response to SRCB immunisation is less dependent on T cell help, and that GC B cells are able to survive and expand for a limited time without T cell help (de Vinuesa et al., 2000). A recent study testing the inhibition of pSYK degradation in GC B cells using mixed bone marrow chimeras (Davidzohn et al., 2020) showed that increased SYK signalling led to an increase in the GC LZ compartment. Further differentiation of these LZ B cells depended on Tfh cell help (Davidzohn et al., 2020; de Vinuesa et al., 2000; Gitlin et al., 2015; Shulman et al., 2013). We show here that SHP-1 deletion in the GC leads to higher levels of pSYK. At least some of these GC B cells are able to recruit efficient Tfh cell help, indicated by the increased expression of cMYC (Calado et al., 2012; Dominguez-Sola et al., 2012). However, many early GC B cells undergo apoptosis, possibly because Tfh cell help is limiting at this early stage. GCs are not only sites of affinity maturation. B cell selection in the GC also guarantees peripheral tolerance (Goodnow et al., 1989; Russell et al., 1991). The data shown here could reflect the deletion of autoreactive GC B cells that encounter inadequate BCR signalling and are not able to recruit adequate Tfh cell help in time. In the same way, higher affinity SHP1-deficient GC B cells may be deleted because they are not recruiting sufficient Tfh cell help. This would indicate that affinity dependent BCR signalling not only is important for affinity dependent B cell selection, but also that the balance of BCR signalling and Tfh cell-mediated rescue may regulate tolerance during the GC B cell responses.

## Author Contributions

Conceptualization, J.C.Y-P. and K.M.T. Investigation, J.C.Y.P., L.Z., R.A.M-A., L.G-I., Y.Z. Formal analysis J.C.Y-P. Resources, Y.A.S., M.S and K.M.T. Writing – Original Draft, J.C.Y-P. Writing – Review & Editing, J.C.Y-P. and K.M.T. All authors reviewed and edited the final version of the manuscript. Supervision and Funding Acquisition, K.M.T.

## Acknowledgments

We are grateful to Benjamin G. Neel (NYU School of Medicine) for Ptpn6^*fl/wt*^ mice and S. Casola (IFOM, Milan, Italy) for Cγ1^*Cre/+*^ mice. We thank Mark J. Shlomchik and Wei Luo for helpful discussion. In addition, we would like to thank at the Biomedical Service Unit, Flow Cytometry Services and Microscopy and Imaging Services at the University of Birmingham. This work was supported by grants from the BBSRC BB/M025292/1 to K.M.T., post-doctoral fellowship program from National Council of Science and Technology-Mexico (CONACYT-Mexico) to J.C.Y-P.

## Conflict of Interest Statement

The authors declare no personal, professional or financial conflict of interest.

## Supplemental material

**Supplementary Figure 1.**
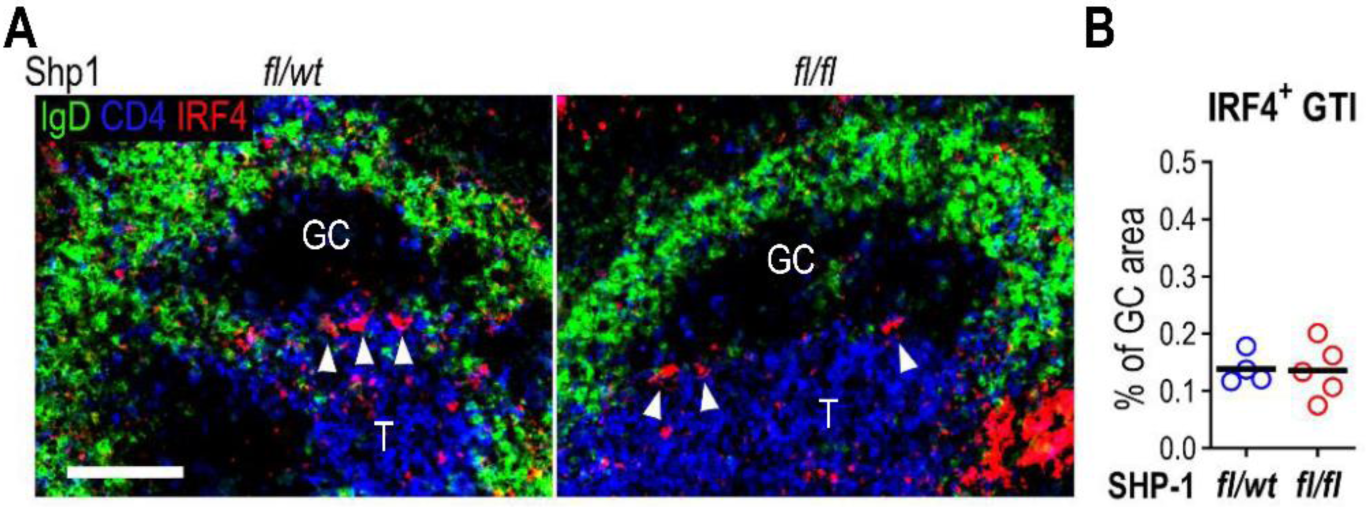
GC-associated PCs during the GC response to SRBCs. Quantification of IRF4^+^ PCs from Cγ1^*Cre/+*^Ptpn6^*fl/wt*^ (*fl/wt*) and Cγ1^*Cre/+*^Ptpn6^*fl/fl*^ (*fl/fl*) mice d5 post SRBC immunisation in the GC-T zone interface (Zhang et al., 2018). Spleens were stained with IRF4 (red), IgD (green), and CD4 (blue). T, T zone; GC, germinal centre. Bar, 100 µm. (% of GC area; results are representative of one of two independent experiments. WT, n=4; Shp1, n=5). Horizontal lines indicate the mean.

**Supplementary Figure 2.**
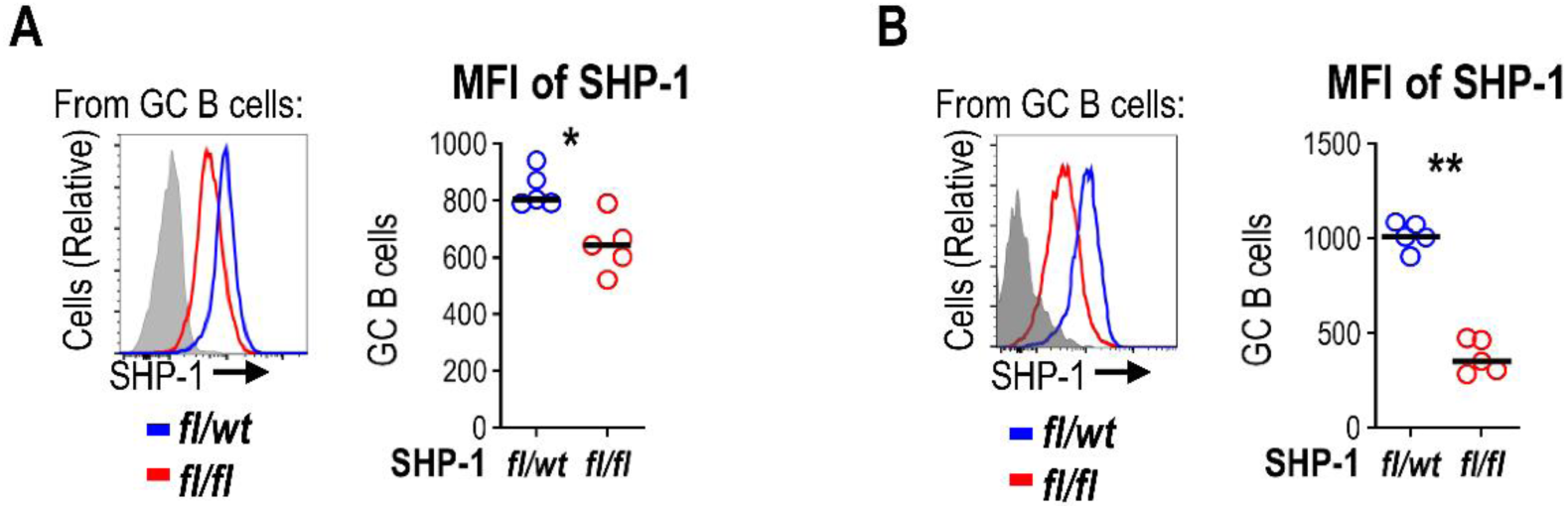
SHP-1 expression is decreased in GC B cells from Cγ1^*Cre/+*^Ptpn6^*fl/fl*^ mice. Cγ1^*Cre/+*^Ptpn6^*fl/wt*^ (*fl/wt*) and Cγ1^*Cre/+*^Ptpn6^*fl/fl*^ (*fl/fl*) mice were immunised with SRBCs (**A**) or NP-CGG (**B**) and SHP-1 expression was determined 5 or 8 days post immunisation, respectively. Representative FACS plot (left) and summary data (right) of SHP-1 expression (MFI of GC B cells; results are representative of one of two independent experiments. WT, n=5; Shp1, n=5); horizontal lines indicate the mean. *P < 0.05, **P < 0.01 (parametric two-tailed unpaired t-test).

**Supplementary Figure 3.**
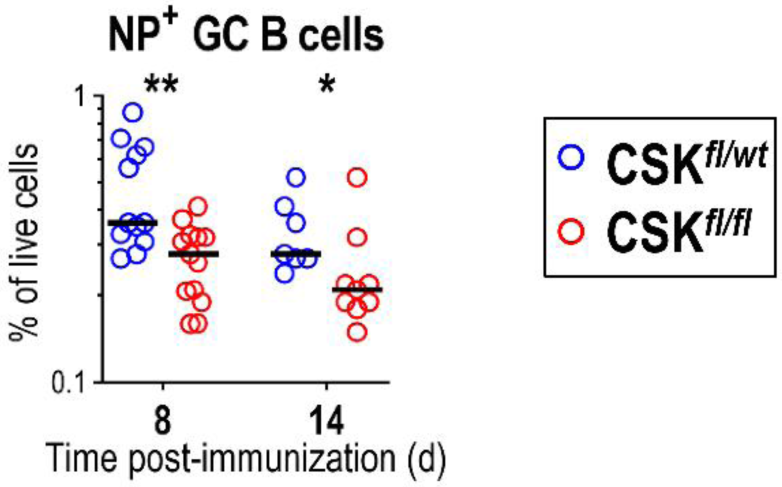
NP specific GC B cells are reduced in the absence of CSK. Cγ1^*Cre/+*^Csk^*fl/wt*^ (open blue circles) and Cγ1^*Cre/+*^Csk^*fl/fl*^ (open red circles) mice were immunised with NP-CGG and GC response was analysed 8 and 14 days post immunisation. (% of live cells; results are combined from two to three independent experiments. WT, n=7-12; Shp1, n=9-13); horizontal lines indicate the mean. *P < 0.05, **P < 0.01 (two-way ANOVA).

## Methods

### RESOURCE AVAILABILITY

#### Lead Contact

Further information and requests for resources and reagents should be directed to and will be fulfilled by Lead Contact, Kai-Michael Toellner.

#### Materials Availability

Materials generated in this study are available upon request.

#### Data and Code Availability

This study did not generate any unique datasets or code.

## EXPERIMENTAL MODEL AND SUBJECT DETAILS

### Mice

Cγ1^Cre/+^ Ptpn6^*fl/wt*^ (Shp1^*fl/wt*^) and Cγ1^*Cre/+*^Ptpn6^*fl/fl*^ (Shp1^*fl/fl*^) mice were generated by the mating of Cγ1^*Cre/+*^ (kindly donated by S Casola, IFOM, Milan, Italy) (Casola et al., 2006) and Ptpn6^*fl/wt*^ animals (Pao et al., 2007). Ptpn6^*fl/wt*^ animals had been backcrossed extensively onto C57BL6. For some experiments, Cγ1^Cre/+^ Ptpn6^*fl/wt*^ mice were crossed with ROSA^mT/mG^ animals (007576; Jackson Laboratory), which contain a Cre-inducible membrane-tagged version of eGFP (Muzumdar et al., 2007). Animal experiments were licensed by the UK Home Office according to the Animals Scientific Procedures Act 1986 and approved by local ethics committee, University of Birmingham, UK.

## METHOD DETAILS

### Immunisation

2 ⨯ 10^8^ sheep red blood cells (SRBCs) (TCS Biosciences, UK) in PBS were injected intravenously in the lateral tail vein. NP (4-hydroxy-3-nitrophenyl acetyl) was conjugated to CGG (Chicken γ-globulin) at a ratio of NP_18_-CGG. Mice were immunised intraperitoneally (i.p.) with 50μg NP_18_-CGG precipitated in alum plus 10^5^ chemically inactivated *Bordetella pertussis* (LEE laboratories, BC, USA) or subcutaneously on the plantar surface of the foot with 20μg NP_18_-CGG precipitated in alum plus 10^5^ chemically inactivated *Bordetella pertussis*.

### Immunofluorescence

Spleens and popliteal lymph nodes obtained at different time-points post-immunisation were frozen and cryosectioned. Slides were rehydrated in PBS and blocked using 1% BSA (Sigma-Aldrich) in PBS for 30 min. Antibodies were diluted at the optimal dilution in PBS, 1% BSA and incubated in a humid dark chamber for 1 h. Allophycocyanin-CD4 (GK1.5), BrilliantViolet421-IgD (11-26c.2a), Alexa633-goat anti-IgG1, goat anti-mouse IRF4 (M-17), rabbit anti-mouse active Caspase 3 (C92-605), and sheep anti-IgD (Abcam) were used. Secondary antibodies were Cy3-conjugated donkey anti-rabbit and Alexa488-conjugated donkey anti-sheep, Alexa555-conjugated donkey anti-goat and streptavidin Alexa555-conjugated. Slides were mounted in ProLong Gold antifade reagent (Invitrogen, UK) and left to dry in a dark chamber for 24 h. Images were taken on an Axio Scan Z1 microscope (Zeiss). Image data were processed using FIJI (Schindelin et al., 2012) or ZEN (Carl Zeiss Germany).

### Flow Cytometry

Red blood cells were or not lysed by ACK lysing buffer (Gibco). Cell suspensions were blocked by CD16/32 (93) diluted in FACS buffer (PBS supplemented with 0.5% BSA plus 2mM EDTA), and then followed with staining cocktail: BrilliantViolet510 B220 (RA3-6B2), BrilliantViolet711 CD138 (281-2), NP-Phycoerythrin for detecting antigen specific B cells (in house), Fluorescein isothiocyanate CD38 (90), BrilliantViolet605 Fas (Jo2), Allophycocyanin IgG1 (X56), Phycoerythrin IgM (Igh-6b), rabbit anti-mouse active caspase 3 (C92-605), streptavidin-Texas Red Phycoerythrin, Phycoerythrin pSYK (I120-722), monoclonal-rabbit anti-SHP-1 (C14H6). Swine anti-rabbit biotin to detect rabbit anti-mouse active caspase 3. Annexin V apoptosis detection kit was used for staining apoptotic and dead cells. For intracellular/intranuclear staining, after surface staining, cell suspensions were treated with the Foxp3/Transcription Factor Fixation/Permeabilization Foxp3 kit (eBioscience, Carlsbad, CA), according to manufacturer specification. Samples were acquired using BD LSRFortessa Analyzer (BD Biosciences, USA) with the software BD FACSDiva (BD Biosciences). Data were analysed with FlowJo v10 (FlowJo LLC, USA).

### ELISA

Serial dilutions of serum samples were analysed by ELISA on NP_15_-BSA (5 μg/ml)–coupled microtiter plates to detect NP-specific IgG1, or NP_2_-BSA (5 μg/ml)–coupled microtiter plates to measure the NP-specific IgM or the high-affinity fraction of IgG1. AP-conjugated secondary antibodies anti-IgM and anti-IgG1 were from Southern Biotech. The substrate of AP was p-nitrophenyl phosphate dissolved in Tris buffer (SIGMAFAST, Sigma-Aldrich). The absorbance was read at 405 nm by using a Synergy HT Microplate Reader (BioTek). OD values were plotted against dilution and smoothed lines were drawn through each dilution series. Relative antibody titres were read as maximal dilution where OD was above an arbitrary threshold. Relative affinity was calculated by dividing ELISA titre derived from NP_2_-BSA– coupled plates by ELISA titre derived from NP_15_-BSA-coupled plates.

### Statistical analysis

All analysis was performed using GraphPad Prism 7 software. To calculate significance parametric two-tailed unpaired t-test and two-way ANOVA were used. Statistics throughout were performed by comparing pooled data obtained from all independent experiments. P values <0.05 were considered significant (*). *p<0.05, ** p< 0.01, *** p<0.001, ****p<0.0001.

## Notes

### Competing Interest Statement

The authors have declared no competing interest.

## References

Adachi, T., Wienands, J., Wakabayashi, C., Yakura, H., Reth, M., and Tsubata, T. (2001). SHP-1 requires inhibitory co-receptors to down-modulate B cell antigen receptor-mediated phosphorylation of cellular substrates. J Biol Chem 276, 26648–26655.

Akkaya, M., Traba, J., Roesler, A.S., Miozzo, P., Akkaya, B., Theall, B.P., Sohn, H., Pena, M., Smelkinson, M., Kabat, J., et al. (2018). Second signals rescue B cells from activation-induced mitochondrial dysfunction and death. Nat Immunol.

Bachmann, M.F., Kundig, T.M., Odermatt, B., Hengartner, H., and Zinkernagel, R.M. (1994). Free recirculation of memory B cells versus antigen-dependent differentiation to antibody-forming cells. J Immunol 153, 3386–3397.

Buchner, M., and Muschen, M. (2014). Targeting the B-cell receptor signaling pathway in B lymphoid malignancies. Curr Opin Hematol 21, 341–349.

Calado, D.P., Sasaki, Y., Godinho, S.A., Pellerin, A., Kochert, K., Sleckman, B.P., de Alboran, I.M., Janz, M., Rodig, S., and Rajewsky, K. (2012). The cell-cycle regulator c-Myc is essential for the formation and maintenance of germinal centers. Nat Immunol 13, 1092–1100.

Cariappa, A., Tang, M., Parng, C., Nebelitskiy, E., Carroll, M., Georgopoulos, K., and Pillai, S. (2001). The follicular versus marginal zone B lymphocyte cell fate decision is regulated by Aiolos, Btk, and CD21. Immunity 14, 603–615.

Casola, S., Cattoretti, G., Uyttersprot, N., Koralov, S.B., Seagal, J., Hao, Z., Waisman, A., Egert, A., Ghitza, D., and Rajewsky, K. (2006). Tracking germinal center B cells expressing germ-line immunoglobulin gamma1 transcripts by conditional gene targeting. Proc Natl Acad Sci U S A 103, 7396–7401.

D’Ambrosio, D., Hippen, K.L., Minskoff, S.A., Mellman, I., Pani, G., Siminovitch, K.A., and Cambier, J.C. (1995). Recruitment and activation of PTP1C in negative regulation of antigen receptor signaling by Fc gamma RIIB1. Science 268, 293–297.

Dal Porto, J.M., Haberman, A.M., Kelsoe, G., and Shlomchik, M.J. (2002). Very low affinity B cells form germinal centers, become memory B cells, and participate in secondary immune responses when higher affinity competition is reduced. J Exp Med 195, 1215–1221.

Davidzohn, N., Biram, A., Stoler-Barak, L., Grenov, A., Dassa, B., and Shulman, Z. (2020). Syk degradation restrains plasma cell formation and promotes zonal transitions in germinal centers. J Exp Med 217.

de Vinuesa, C.G., Cook, M.C., Ball, J., Drew, M., Sunners, Y., Cascalho, M., Wabl, M., Klaus, G.G., and MacLennan, I.C. (2000). Germinal centers without T cells. J Exp Med 191, 485–494.

Dominguez-Sola, D., Victora, G.D., Ying, C.Y., Phan, R.T., Saito, M., Nussenzweig, M.C., and Dalla-Favera, R. (2012). The proto-oncogene MYC is required for selection in the germinal center and cyclic reentry. Nat Immunol 13, 1083–1091.

Gitlin, A.D., Mayer, C.T., Oliveira, T.Y., Shulman, Z., Jones, M.J., Koren, A., and Nussenzweig, M.C. (2015). HUMORAL IMMUNITY. T cell help controls the speed of the cell cycle in germinal center B cells. Science 349, 643–646.

Goodnow, C.C., Crosbie, J., Jorgensen, H., Brink, R.A., and Basten, A. (1989). Induction of self-tolerance in mature peripheral B lymphocytes. Nature 342, 385–391.

Hanayama, R., Tanaka, M., Miyasaka, K., Aozasa, K., Koike, M., Uchiyama, Y., and Nagata, S. (2004). Autoimmune disease and impaired uptake of apoptotic cells in MFG-E8-deficient mice. Science 304, 1147–1150.

Khalil, A.M., Cambier, J.C., and Shlomchik, M.J. (2012). B cell receptor signal transduction in the GC is short-circuited by high phosphatase activity. Science 336, 1178–1181.

Kouskoff, V., Famiglietti, S., Lacaud, G., Lang, P., Rider, J.E., Kay, B.K., Cambier, J.C., and Nemazee, D. (1998). Antigens varying in affinity for the B cell receptor induce differential B lymphocyte responses. J Exp Med 188, 1453–1464.

Kraus, M., Alimzhanov, M.B., Rajewsky, N., and Rajewsky, K. (2004). Survival of resting mature B lymphocytes depends on BCR signaling via the Igalpha/beta heterodimer. Cell 117, 787–800.

Lam, K.P., Kuhn, R., and Rajewsky, K. (1997). In vivo ablation of surface immunoglobulin on mature B cells by inducible gene targeting results in rapid cell death. Cell 90, 1073–1083.

Li, Y.F., Xu, S., Ou, X., and Lam, K.P. (2014). Shp1 signalling is required to establish the long-lived bone marrow plasma cell pool. Nat Commun 5, 4273.

Luo, W., Weisel, F., and Shlomchik, M.J. (2018). B Cell Receptor and CD40 Signaling Are Rewired for Synergistic Induction of the c-Myc Transcription Factor in Germinal Center B Cells. Immunity 48, 313–326 e315.

MacLennan, I.C. (1994). Germinal centers. Annu Rev Immunol 12, 117–139.

MacLennan, I.C., Toellner, K.M., Cunningham, A.F., Serre, K., Sze, D.M., Zuniga, E., Cook, M.C., and Vinuesa, C.G. (2003). Extrafollicular antibody responses. Immunol Rev 194, 8–18.

Maeda, A., Kurosaki, M., Ono, M., Takai, T., and Kurosaki, T. (1998). Requirement of SH2-containing protein tyrosine phosphatases SHP-1 and SHP-2 for paired immunoglobulin-like receptor B (PIR-B)-mediated inhibitory signal. J Exp Med 187, 1355–1360.

Marshall, J.L., Zhang, Y., Pallan, L., Hsu, M.C., Khan, M., Cunningham, A.F., MacLennan, I.C., and Toellner, K.M. (2011). Early B blasts acquire a capacity for Ig class switch recombination that is lost as they become plasmablasts. Eur J Immunol 41, 3506–3512.

Mueller, J., Matloubian, M., and Zikherman, J. (2015). Cutting edge: An in vivo reporter reveals active B cell receptor signaling in the germinal center. J Immunol 194, 2993–2997.

Muzumdar, M.D., Tasic, B., Miyamichi, K., Li, L., and Luo, L. (2007). A global double-fluorescent Cre reporter mouse. Genesis 45, 593–605.

Nguyen, T.T., Klasener, K., Zurn, C., Castillo, P.A., Brust-Mascher, I., Imai, D.M., Bevins, C.L., Reardon, C., Reth, M., and Baumgarth, N. (2017). The IgM receptor FcmuR limits tonic BCR signaling by regulating expression of the IgM BCR. Nat Immunol 18, 321–333.

Niiro, H., and Clark, E.A. (2002). Regulation of B-cell fate by antigen-receptor signals. Nat Rev Immunol 2, 945–956.

Nitschke, L., and Tsubata, T. (2004). Molecular interactions regulate BCR signal inhibition by CD22 and CD72. Trends Immunol 25, 543–550.

O’Connor, B.P., Vogel, L.A., Zhang, W., Loo, W., Shnider, D., Lind, E.F., Ratliff, M., Noelle, R.J., and Erickson, L.D. (2006). Imprinting the fate of antigen-reactive B cells through the affinity of the B cell receptor. J Immunol 177, 7723–7732.

Oropallo, M.A., and Cerutti, A. (2014). Germinal center reaction: antigen affinity and presentation explain it all. Trends Immunol 35, 287–289.

Pao, L.I., Lam, K.P., Henderson, J.M., Kutok, J.L., Alimzhanov, M., Nitschke, L., Thomas, M.L., Neel, B.G., and Rajewsky, K. (2007). B cell-specific deletion of protein-tyrosine phosphatase Shp1 promotes B-1a cell development and causes systemic autoimmunity. Immunity 27, 35–48.

Parry, S.L., Hasbold, J., Holman, M., and Klaus, G.G. (1994). Hypercross-linking surface IgM or IgD receptors on mature B cells induces apoptosis that is reversed by costimulation with IL-4 and anti-CD40. J Immunol 152, 2821–2829.

Paus, D., Phan, T.G., Chan, T.D., Gardam, S., Basten, A., and Brink, R. (2006). Antigen recognition strength regulates the choice between extrafollicular plasma cell and germinal center B cell differentiation. J Exp Med 203, 1081–1091.

Roco, J.A., Mesin, L., Binder, S.C., Nefzger, C., Gonzalez-Figueroa, P., Canete, P.F., Ellyard, J., Shen, Q., Robert, P.A., Cappello, J., et al. (2019). Class-Switch Recombination Occurs Infrequently in Germinal Centers. Immunity 51, 337–350 e337.

Russell, D.M., Dembic, Z., Morahan, G., Miller, J.F., Burki, K., and Nemazee, D. (1991). Peripheral deletion of self-reactive B cells. Nature 354, 308–311.

Schindelin, J., Arganda-Carreras, I., Frise, E., Kaynig, V., Longair, M., Pietzsch, T., Preibisch, S., Rueden, C., Saalfeld, S., Schmid, B., et al. (2012). Fiji: an open-source platform for biological-image analysis. Nat Methods 9, 676–682.

Schwickert, T.A., Victora, G.D., Fooksman, D.R., Kamphorst, A.O., Mugnier, M.R., Gitlin, A.D., Dustin, M.L., and Nussenzweig, M.C. (2011). A dynamic T cell-limited checkpoint regulates affinity-dependent B cell entry into the germinal center. J Exp Med 208, 1243–1252.

Shulman, Z., Gitlin, A.D., Targ, S., Jankovic, M., Pasqual, G., Nussenzweig, M.C., and Victora, G.D. (2013). T follicular helper cell dynamics in germinal centers. Science 341, 673–677.

Steinhoff, U., Muller, U., Schertler, A., Hengartner, H., Aguet, M., and Zinkernagel, R.M. (1995). Antiviral protection by vesicular stomatitis virus-specific antibodies in alpha/beta interferon receptor-deficient mice. J Virol 69, 2153–2158.

Stewart, I., Radtke, D., Phillips, B., McGowan, S.J., and Bannard, O. (2018). Germinal Center B Cells Replace Their Antigen Receptors in Dark Zones and Fail Light Zone Entry when Immunoglobulin Gene Mutations are Damaging. Immunity 49, 477–489 e477.

Toellner, K.M., Luther, S.A., Sze, D.M., Choy, R.K., Taylor, D.R., MacLennan, I.C., and Acha-Orbea, H. (1998). T helper 1 (Th1) and Th2 characteristics start to develop during T cell priming and are associated with an immediate ability to induce immunoglobulin class switching. J Exp Med 187, 1193–1204.

Toellner, K.M., Sze, D.M., and Zhang, Y. (2018). What Are the Primary Limitations in B-Cell Affinity Maturation, and How Much Affinity Maturation Can We Drive with Vaccination? A Role for Antibody Feedback. Cold Spring Harb Perspect Biol 10.

Tsiantoulas, D., Kiss, M., Bartolini-Gritti, B., Bergthaler, A., Mallat, Z., Jumaa, H., and Binder, C.J. (2017). Secreted IgM deficiency leads to increased BCR signaling that results in abnormal splenic B cell development. Sci Rep 7, 3540.

Tsubata, T., Murakami, M., and Honjo, T. (1994a). Antigen-receptor cross-linking induces peritoneal B-cell apoptosis in normal but not autoimmunity-prone mice. Curr Biol 4, 8–17.

Tsubata, T., Murakami, M., Nisitani, S., and Honjo, T. (1994b). Molecular mechanisms for B lymphocyte selection: induction and regulation of antigen-receptor-mediated apoptosis of mature B cells in normal mice and their defect in autoimmunity-prone mice. Philos Trans R Soc Lond B Biol Sci 345, 297–301.

Victora, G.D., and Nussenzweig, M.C. (2012). Germinal centers. Annu Rev Immunol 30, 429–457.

Victora, G.D., Schwickert, T.A., Fooksman, D.R., Kamphorst, A.O., Meyer-Hermann, M., Dustin, M.L., and Nussenzweig, M.C. (2010). Germinal center dynamics revealed by multiphoton microscopy with a photoactivatable fluorescent reporter. Cell 143, 592–605.

Watanabe, N., Nomura, T., Takai, T., Chiba, T., Honjo, T., and Tsubata, T. (1998). Antigen receptor cross-linking by anti-immunoglobulin antibodies coupled to cell surface membrane induces rapid apoptosis of normal spleen B cells. Scand J Immunol 47, 541–547.

Weisel, F.J., Zuccarino-Catania, G.V., Chikina, M., and Shlomchik, M.J. (2016). A Temporal Switch in the Germinal Center Determines Differential Output of Memory B and Plasma Cells. Immunity 44, 116–130.

Yam-Puc, J.C., Zhang, L., Zhang, Y., and Toellner, K.M. (2018). Role of B-cell receptors for B-cell development and antigen-induced differentiation. F1000Res 7, 429.

Zhang, Y., Garcia-Ibanez, L., and Toellner, K.M. (2016). Regulation of germinal center B-cell differentiation. Immunol Rev 270, 8–19.

Zhang, Y., Tech, L., George, L.A., Acs, A., Durrett, R.E., Hess, H., Walker, L.S.K., Tarlinton, D.M., Fletcher, A.L., Hauser, A.E., et al. (2018). Plasma cell output from germinal centers is regulated by signals from Tfh and stromal cells. J Exp Med 215, 1227–1243.

